# Empowering High Throughput Screening of 3D Models: Automated Dispensing of Cervical and Endometrial Cancer Cells

**DOI:** 10.1101/2024.03.06.583783

**Authors:** Samantha Seymour, Ines Cadena, Mackenzie Johnson, Molly Jenne, Iman Adem, Alyssa Almer, Rachel Frankovic, Danielle Spence, Andrea Haddadin, Kaitlin C. Fogg

**Affiliations:** School of Chemical, Biological, and Environmental Engineering, Oregon State University, Corvallis, OR; Lonza, Bend, OR

## Abstract

**Purpose:** Cervical and endometrial cancers pose significant challenges in women’s healthcare due to their high mortality rates and limited treatment options. High throughput screening (HTS) of cervical and endometrial cancer *in vitro* models offer a promising avenue for drug repurposing and broadening patient treatment options. Traditional two-dimensional (2D) cell-based screenings have limited capabilities to capture crucial multicellular interactions, that are improved upon in three dimensional (3D) multicellular tissue engineered models. However, manual fabrication of the 3D platforms is both time consuming and subject to variability. Thus, the goal of this study was to utilize automated cell dispensing to fabricate 3D cell-based HTS platforms using the HP D100 Single Cell Dispenser to dispense cervical and endometrial cancer cells.

**Methods:** We evaluated the effects of automated dispensing of the cancer cell lines by tuning the dispensing protocol to align with cell size measured in solution and the minimum cell number for acceptable cell viability and proliferation. We modified our previously reported coculture models of cervical and endometrial cancer to be in a 384 well plate format and measured microvessel length and cancer cell invasion.

**Results:** Automatically and manually dispensed cells were directly compared revealing minimal differences between the dispensing methods. These findings suggest that automated dispensing of cancer cells minimally affects cell behavior and can be deployed to decrease *in vitro* model fabrication time.

**Conclusions:** By streamlining the manufacturing process, automated dispensing holds promise for enhancing efficiency and scalability of 3D *in vitro* HTS platforms, ultimately contributing to advancement in cancer research and treatment.

## 1. Introduction

Endometrial and cervical cancer are two of the most common cancers that led to over 439,000 deaths in 2020 globally.^1,2^ In the same year, over 417,000 women were diagnosed with endometrial cancer and over 604,000 women were diagnosed with cervical cancer.^1,2^ Treatment for these diseases is unique for each patient and varies based on disease progression, often entailing a combination of surgery, radiotherapy, and chemotherapy. Radical hysterectomy surgery has side effects such as early menopause, excessive bleeding, infections, nerve injury, and vaginal cuff dehiscence, a rare but possible complication where the abdominal contacts are expelled through the vaginal opening.^3,4^ Combination therapy such as chemotherapy before surgery can help reduce the tumor size and extent of surgery, and for cervical cancer patients has been shown to decreased recurrence rates compared to chemotherapy treatments.^5,6^ Doxorubicin, paclitaxel, cisplatin, and carboplatin are among the most common single-agent chemotherapy treatments, but higher response rates have been observed for cervical cancer patients using combination chemotherapy treatments such as carboplatin-paclitaxel or cisplatin-paclitaxel.^5,7^ However, chemotherapy can have significant side effects that include: tiredness, sickness, hair loss, anemia, sore mouth, loss of appetite, memory and concentration problems, reduced fertility, and emotional distress to list a few.^8^ Not only do the current treatment methods have significant side effects, but only a fraction of patients are responsive to the available treatment methods. Thus, there is a clear need for new therapy options to increase the quality of treatment and patient response rate. However, bringing novel therapeutics to market remains a challenge and only 7% of new anticancer drugs are approved for clinical usage.^9^ Alternatively, instead of creating novel therapeutics, drug repurposing can provide another avenue for increasing treatment options. There are over 19,000 prescription drugs already approved by the FDA with known safety profiles.^10–12^ By identifying which of these familiar drugs have anticancer properties, we can quickly expand the treatment options available to patients.

One approach for drug repurposing is high throughput screening (HTS). Cell-based HTS is focused on cell behavior in the presence or absence of a candidate compound.^12^ Two dimensional (2D) cell-based HTS can inform biochemical and physical properties effected by the candidate compound by measuring cell adhesion, proliferation, and differentiation.^13^ However, the 2D cell-based HTS only provides information of the single cell response and lacks the ability to capture cell to cell interactions, or cell phenotypic responses characteristic of the candidate tissue.^13^ In context, three dimensional (3D) cell-based HTS assays that resemble the tissue microenvironment can demonstrate more complex and dynamic cell interactions.^13–15^ However, scaling up these complex *in vitro* screens is limited by the lab techniques required to build the models and operate the HTS platforms.^16–18^

To integrate 3D *in vitro* models with HTS for drug repurposing, it is necessary to make the models scalable for drug library screening. Currently there are over 19,000 drugs approved by the U.S. Food & Drug Association (FDA) that could be screened for anticancer behaviors such as preventing microvessel formation and cancer invasion.^10^ Screening just one dose of each product in the FDA drug library would require 208 96-well plates or 52 384 well plates. Besides the large number of plates required to explore the FDA approved library, the fabrication of 3D models manually is time consuming and be variable between each model.^19,20^ Depending on the complexity of the model, manually fabricating just a few 96 well plates can take two researchers the majority of a day to complete. The fabrication of these models can be scaled to reach drug screening goals by utilizing automatic cell dispensing to decrease the time and variability between wells. Automated cell dispensing comes in two main forms: single cell dispensing and bulk cell dispensing. Single cell dispensing excels with dispensing a limited number of cells precisely, while bulk cell dispensing can achieve high density seeding.^21,22^ Depending on the number of cells required for the *in vitro* model, a single cell or bulk cell dispenser can be utilized to dispense the cells in an efficient and reproducible manner.

In this paper, our objective was to automate cancer cell dispensing of our previously reported *in vitro* HTS multilayer multicellular models for endometrial and cervical cancers.^23,24^ We evaluated the effects of automated dispensing of the cancer cell lines by tuning the dispensing protocol to align with cell size measured in solution and the minimum cell number for acceptable cell viability and proliferation. We modified our previously reported coculture models of cervical and endometrial cancer to be in a 384 well plate format and measured microvessel length and cancer cell invasion. Overall, this study demonstrates that there are few significant differences in the phenotypic responses when dispensing the cancer cell lines are dispensed automatically compared to manually in the *in vitro* model, opening future avenues to automatically manufacture more multilayer multicellular models for future HTS drug repurposing investigations.

## 2. Materials and Methods

### 2.1 Cell Lines and Reagents

Unless otherwise stated, all reagents were purchased from ThermoFisher (Waltham, MA). Human microvascular endothelial cells (hMVEC) and human umbilical vein endothelial cells (hUVEC) were obtained from Lonza (hMVEC CC-2543, hUVEC C2519A, Walkersville, MD) and used without additional characterization. The hMVEC cells were expanded in EGM-2 MV media (EBM-2 (CC – 3203) supplemented with Lonza’s SingleQuot supplements (CC – 4147), Lonza) and further supplemented with 5% fetal bovine serum (FBS) until use at passage 5. The hUVEC cells were cultured and expanded until passage 5 with EGM-2 media (CC-3156, Lonza) and SingleQuot supplements (CC-4176, Lonza). Human cervical cancer cell lines SiHa (ATCC^®^ HTB-35™), and Ca Ski (ATCC CRM-CRL-1550) and human endometrial cancer cell line HEC-1A (ATCC HTB-112™), were purchased from ATCC (Manassas, VA) and used without additional characterization. The cancer cell lines were cultured in Eagle’s Minimum Essential Medium (EMEM, ATCC), RPMI-1640 Medium (ATCC), or McCoy’s 5A medium (ATCC) supplemented by 1% penicillin–streptomycin (Sigma-Aldrich, St. Louis, MO, USA) and 10% FBS, until use at passage 5. All cell types were expanded in standard cell culture conditions (37°C, 21% O_2_, 5% CO_2_) and subcultured before they reached 80% confluency. (**Table 1**)

**Table 1.** Gynecological cancer and endothelial cell lines materials and handling information.

| **Cell Type** | **Passage Used** | **Type** | **Tissue Location** | **Supplier** | **Cell Culture Media** | **Cell Culture Media & Supplier Information** |
| --- | --- | --- | --- | --- | --- | --- |
| **SiHa**<br>**(ATCC® HTB-35™)** | 5 | Grade II, Squamous Cell Carcinoma | Cervix | ATCC<br>Manassas, VA | EMEM | 89% Eagle's Minimum Essential Medium (EMEM) (ATCC 30-2003), 10% FBS, 1% Penicillin - Streptomycin (P/S, Sigma-Aldrich) |
| **Ca Ski**<br>**(ATCC CRM-CRL-5)** | 5 | Epidermoid Carcinoma | Cervix | ATCC<br>Manassas, VA | RPMI-1640 | 89% RPMI (ATCC 30-2001), 10% FBS, 1% P/S |
| **HEC-1A**<br>**(ATCC HTB-112™)** | 5 | Adenocarcinoma | Uterus, Endometrium | ATCC<br>Manassas, VA | McCoy's 5A | 89% McCoy's 5A Medium Modified (ATCC 30-2007), 10% FBS, 1% P/S |
| **hMVEC-DAd**<br>**(Lonza CC-2543)** | 5 | Adult | Dermal Microvascular Endothelial | Lonza,<br>Walkersville, MD | EGM-2MV BulletKit™ Medium (Lonza CC-3202) | 95% EBM-2 + SingleQout supplements (hydrocortisone, human basic fibroblast growth factor (FGF2), human vascular endothelial growth factor (VEGF), human insulin-like growth factor (IGF), human epidermal growth factor (hEGF), ascorbic acid, gentamycin) (Lonza CC-4147), 5% FBS |
| **hUVEC**<br>**(Lonza C2519A)** | 5 | Pooled Donors | Umbilical Vein Endothelial | Lonza,<br>Walkersville, MD | EGM-2 BulletKit™ Medium (Lonza CC-3156) | 95% EBM-2 + SingleQout supplements (hydrocortisone, human basic fibroblast growth factor (FGF2), human vascular endothelial growth factor (VEGF), human insulin-like growth factor (IGF), human epidermal growth factor (hEGF), ascorbic acid, gentamycin, heparin) (Lonza CC-4146), 5% FBS |

Cell size was determined for cancer cell lines (SiHa, Ca Ski, HEC-1A) and endothelial cell lines (hMVEC, hUVEC) either in suspension or adhered to tissue culture plastic (TCP). The diameter of cells adhered to TCP was measured by the Cytation 5 cell imaging multimode reader measurement tool (Agilent Technologies, Santa Clara, CA). In-suspension measurements were determined using the T4 Auto Cellometer Bright Field Cell Counter (Nexelom Bioscience, Lawrence, MA). A well-mixed sample of 20 *μ*L was used to create a 1:1 mixture of cell suspension: Trypan Blue (0.2% w/v diluted with 1X phosphate buffered saline (PBS)). This mixture was pipetted into the measuring chamber, and the total mean cell diameter was recorded.

### 2.2 Cell Dispensing

Dispensing performed by hand pipetting will be referred to as manual dispensing, while automatic dispensing refers to reagents or cell suspensions that were aliquoted by the HP D100 Single Cell Dispenser (HP, Corvallis, OR). The HP D100 Dispensing software, version 3.5.0, was used for all automatic dispensing. The automatic dispenser was prepared and calibrated as specified by the manufacturer. All automatic dispensing was completed using the 96, 384, or custom well plate types. HP specialists assisted in creating the custom wells that allowed 30 cells to be automatically dispensed within a single well. A dispensing grid was created in a 5×6 unit pattern with 0.25 mm between units. Custom wells were created for the center wells of row F within a 384-well plate.

In preparation for dispense, the fluid type was selected to reflect the cell size as measured in suspension. One cell was designated per well of a 96-well plate or per unit in the custom wells, and plate shaking was deactivated. The desired plate was loaded into the carrier, and the cassette was secured in the dispensing head. The cells were prepared by suspension in 1X PBS at 10,000 cells/mL and filtered with a 35 µm filter tube. The suspension was then loaded into the cassette reservoir as recommended by the dispensing software. A deionizer was deployed for five seconds before running the dispensing protocol. During dispense, the suspension travels through a microfluidic channel to a pinch point (funnel) (**Fig. 1**). This is a constriction that detects cells passing through the channel. The pinch point diameter is variable based on the cassette type and should be aligned with the diameter of the cells as recommended by the manufacturer. Immediately after dispensing, the plate was returned to the incubator or tissue culture hood for further preparation. All automatic dispensing was performed on the bench top, and manual dispensing was performed in a tissue culture hood.

**Figure 1.**
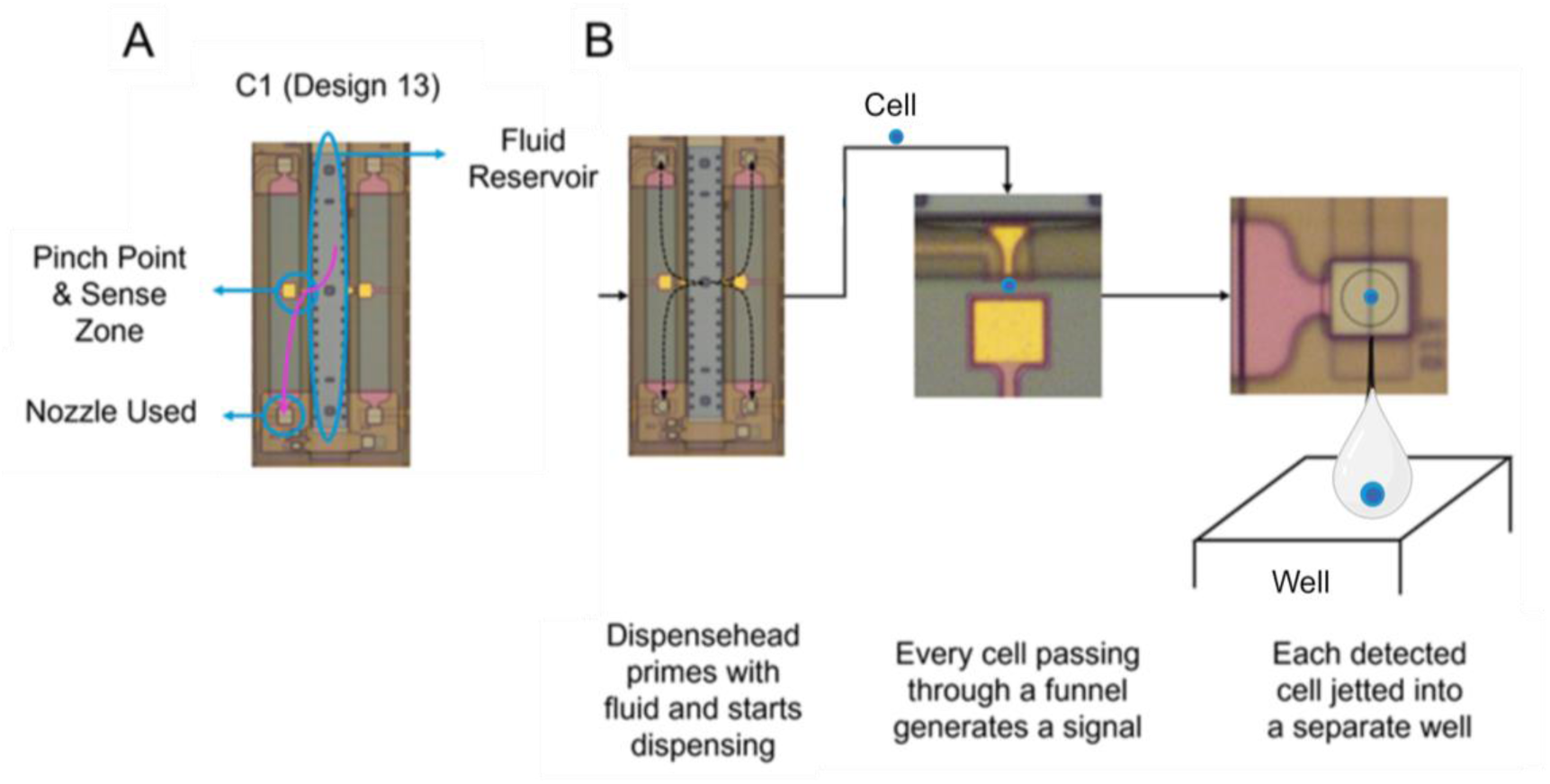
Diagram of dispensing with HP D100 Single Cell Dispenser Cassette, C1 Design 13 (A) and dispensing flow path (B).

Wells predicted and observed to be single cell wells were described as a true positive (TP) result, while wells predicted and observed to be non-single-cell wells represented a true negative (TN) result. However, wells predicted to be single cell wells but observed to be non-single cell wells were indicative of a false positive (FP), and wells predicted to be non-single cell wells but observed to be single cell wells were a false negative (FN) result. This data was then used to determine the true positive rate (TPR) or the rate of wells that were correctly classified by the dispenser (**Equation 1**), and the false positive rate (FPR) or the rate of wells wrongly classified as single cell wells (**Equation 2**):

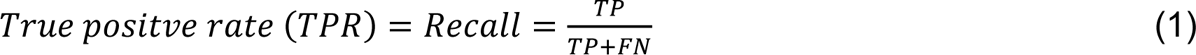

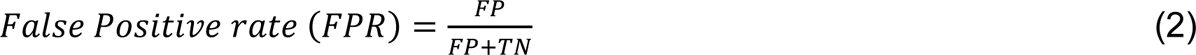

### 2.3 Cell Response

Cell viability and cell number were determined by fluorescent labeling with Calcein AM (2 *μ*M, Calcein AM Viability Dye, 65-0853-39) and Propidium Iodide (5 *μ*M Propidium Iodide Ready Flow™ Reagent, R37169). Cells were rinsed with 2 mL of 1X PBS, and incubated in 2 mL of Calcein AM Propidium Iodide solution for 15 minutes. This was done for both suspended and adherent cells according to manufacturer instructions. Cell nuclei were fluorescently labeled with Hoestch (0.0005 mM, Hoechst 33342 Solution) according to manufacturer instructions. Cell viability was determined by the following equation, (**Equation 3**):

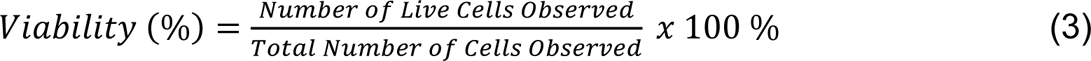

Cell number was used to determine proliferation rate and precision as defined in **Equations 4** and **5**, respectively:

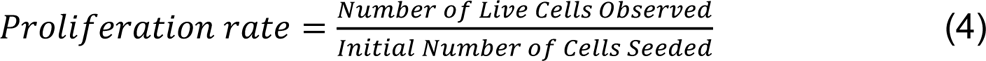

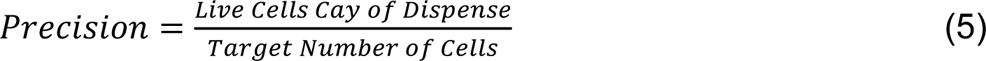

To visualize cells in co-culture, cancer cells and endothelial cells were labeled with CellTracker Red (CellTracker™ Red CMTPX Dye) and CellTracker Green (CellTracker™ Green CMFDA Dye), respectively, as recommend by the manufacturer. Fluorescence was captured by the Cytation 5 cell imaging multimode reader. The phenotypic responses: cancer cell displacement, number of cancer cells per well, endothelial cell coverage, and microvessel length were determined as previously reported.^28^ Briefly, fluorescent images were evaluated by Fiji-ImageJ (NIH, Bethesda, MD) to determine the phenotypic responses. Cancer cell displacement was defined as the farthest point of cancer cell displacement from the initial height after 24 hours of co-culture. The number of cancer cells per well were measured by counting the number of visible cells 48 hours after culture. Cell coverage was measured by calculating the area within each well that is covered by cells and dividing it by the total well area. The average length of endothelial microvessels were measured using FIJI.

### 2.4 Hydrogel Fabrication

Gelatin methacryloyl (GelMA, 7% w/v, Advanced BioMatrix, San Diego, CA) was crosslinked using the UV photo crosslinker Irgacure 2959 (BioMatrix) for 1 minute under a 365 nm UV light crosslinking source according to protocols specified by the manufacturer. Growth factor deficient Matrigel basement membrane matrix (9.2 mg/mL protein concentration, Corning, MA, USA) was used as a control and prepared according to protocols specified by the manufacturer.

The cells of co-culture experiments were suspended in a multilayer multicellular model as previously described.^28^ Briefly, the hydrogel layers were prepared using the concentrations described in **Table 2**. For the cervical cancer model, the bottom hydrogel layer consisted of a combination of collagen type 1 (Col1) (FibriCol, BioMatrix), fibrinogen (FG) from human plasma (30 mg/mL, Sigma-Aldrich), thrombin from human plasma (Sigma-Aldrich) and GelMA (8.7% w/v), and the top hydrogel layer consisted of Col1, fibronectin (FN) (BioMatrix), and GelMA (8.7% w/v). The top layer of the endometrial cancer model consisted of Collagen type IV (ColIV, BioMatrix), fibronectin, human laminin (Sigma-Aldrich) and polyethylene glycol diacrylate (PEGDA, 10% w/v, BioMatrix) crosslinked using the UV photo crosslinker Irgacure 2959 for 1 minute under a 365 nm UV light crosslinking source. For the hydrogels, the pH was adjusted by adding 0.1 N NaOH and PBS to achieve a target pH of seven. (**Table 2**)

**Table 2.** Cervical cancer (CC) and endometrial cancer (EC) hydrogel compositions.

| Model | Layer | Col I (mg/mL) | Col IV (mg/mL) | FG (mg/mL) | FN (mg/mL) | GelMA (%) | Laminin (µg/mL) | PEGDA (%) |
| --- | --- | --- | --- | --- | --- | --- | --- | --- |
| Cervical Cancer | Top | 1.12 | - | - | 0.16 | 8.7 | - | - |
|  | Bottom | 2.5 | - | 2.5 | - | 2.5 | - | - |
| Endometrial Cancer | Top | - | 0.13 | - | 0.17 | - | 0.5 | 11.7 |
|  | Bottom | 1.91 | - | 0.6 | - | 8.7 | - | - |

### 2.5 Monoculture Hydrogel Fabrication

Cancer cell lines (SiHa, Ca Ski, HEC-1A) were seeded on TCP, GelMA (89% 1X PBS 7% GelMA and 1% Irgacure), and Matrigel platforms. Wells containing gels (GelMA or Matrigel) were prepared by manually dispensing 10 μL of the pre-gel mixture into each well. After polymerizing the pre-gel mixtures, 25 μL of cell media was manually dispensed in each well. Cells were either manually or automatically dispensed into the wells. Cells manually dispensed for the purpose of determining the minimum number of cancer cells required per well, were seeded on a decreasing half-log scale starting at 1750 cells per well to 7 cells per well according to **Table 3** (n=3). Cell viability and cell number were determined on days 0 and 7. The cell culture medium was refreshed every other day.

**Table 3.** Seeding densities and dispensing volumes for minimum occupancy investigation.

| Number of Cells Dispensed per Well | 384 Well Plate Column | Cell Density (Cells/mL) | Cell Suspension Volume (μL) |
| --- | --- | --- | --- |
| 1750 | 1 | 100,000 | 17.50 |
| 875 | 2 | 100,000 | 8.75 |
| 438 | 3 | 100,000 | 4.38 |
| 218 | 4 | 10,000 | 21.88 |
| 109 | 5 | 10,000 | 10.92 |
| 55 | 6 | 10,000 | 5.47 |
| 27 | 7 | 1,000 | 27.34 |
| 14 | 8 | 1,000 | 13.67 |
| 7 | 9 | 1,000 | 6.84 |

### 2.6 Multilayer Hydrogel Fabrication

The multilayer multicellular models were fabricated in 384 well plates by layering 10 µL of the bottom hydrogel formulation into each well. Then CellTracker Green labeled hMVEC cells in EGM-2 MV media were pipetted on top of each gel at 0.5 cells/mL in 40 µL. The endothelial cells were allowed to attach for four hours at 37°C and 5% CO_2_. Carefully the media was then removed, and 25 µL of the top hydrogel formulation was pipetted on top of the hMVEC cells. After the hydrogel was crosslinked, CellTracker Red labeled cancer cells were either manually or automatically dispensed on top of the hydrogel to achieve 30 cells per well. To prepare for cell dispense, 10 µL of media were pipetted into each well, the media composed of a 1:1 ratio of EGM-2 MV and cancer cells medium. Multilayer Matrigel constructs were used as a controls, and made using the same methods described above, except the hydrogel formulations were replaced with growth factor deficient Matrigel basement membrane matrix in both layers. The media was changed every 24 hours, and the plates were incubated at 37°C and 5% CO_2_.

### 2.7 Statistical Analysis

All statistical analyses were performed using Prism 9.4.1 software (GraphPad, San Diego, CA). All *p-values* less than 0.05 were considered statistically significant.

## Results

### 3.1 Minimum Occupancy

To determine the minimum number of cancer cells per well in a 384 well plate that could be dispensed and still have acceptable cell viability and proliferation rates, we examined a range from 7 to 1750 cells per well. The success criteria were average viability greater than 50% and average proliferation greater than one-fold change from day seven to day one.

For cells seeded on TCP, SiHa and Ca Ski exceed the success criteria of 50% for all seeding densities (**Fig. 2A**). However, HEC-1A required a minimum of 27 cells per well to achieve the same criteria, with viability generally increasing as cell number increased. For all seeding densities SiHa and Ca Ski achieved the proliferation success criteria, while HEC-1A required 27 cells per well (**Fig. 2B**). Maximum proliferation for SiHa and Ca Ski occurred at 55 and 27 cells per well, respectively. HEC-1A proliferated at a seeding density of 27 cells per well, peaking at 55 cells per well. Ca Ski and HEC-1A cell lines observed a normal distribution trend, while SiHa observed positive skewness (**Fig. 2B**). For TCP, all cell lines observed increased viability as the number of cells per well increased, while the proliferation rate trends reached a peak value and decreased as the number of cells increased. For cells seeded on GelMA, SiHa and HEC-1A were viable for all seeding densities with increasing cell viability as the number of cells per well increased (**Fig. 2C**). However, Ca Ski only achieved average viability greater than 50% at 7 cells per well. All three cell lines met our success criteria for proliferation rate, but they did not proliferate well on GelMA (**Fig. 2D**). Lastly, SiHa seeded on Matrigel required at least 27 cells per well, Ca Ski did not meet the success criteria for any cell densities, and HEC-1A met the criteria for all seeding densities (**Fig. 2E**). All three cell lines exhibited decreasing proliferation with increased cells per well, with 14 cells per well meeting the success criteria for all three cell lines on Matrigel (**Fig. 2F**). Moving forward these data informed us that we needed a minimum of 27 cells per well to observe viable and proliferating cells per well for all platforms in 384 well plates.

**Figure 2.**
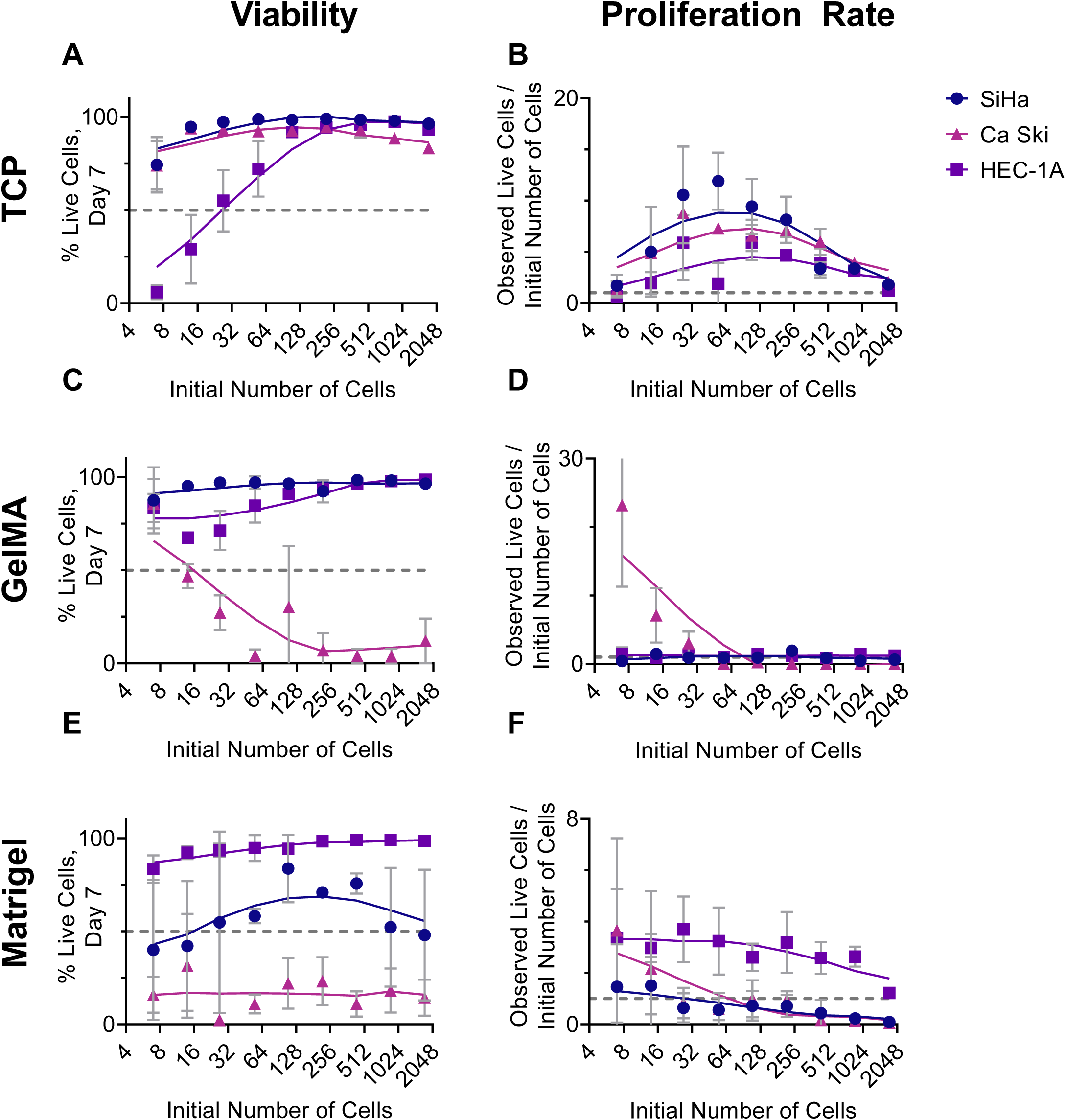
Comparison of viability (%) on the platforms: TCP (**A**), GelMA (**C**), Matrigel (**E**), and proliferation rate: TCP (**B**), GelMA (**D**), Matrigel (**F**), for the cancer cell lines (SiHa, Ca Ski, HEC-1A). Cells were manually dispensed on top of 25 μL of cell specific culture medium on TCP, 10 μL of GelMA, or 10 μL Matrigel in a 384 well plate. Data represents the mean ± SD (n=3).

### 3.2 Pinch point Diameter

To choose between an 11 µm and 14 µm pinch point diameter for dispensing cells with the HP D100 Single Cell Dispenser we first recoded the diameter of the cell lines in suspension and on TCP (**Fig. 3**). Secondly, we evaluated the true positive and true negative rates for dispensing. (**Sup. Table 2**)

**Figure 3.**
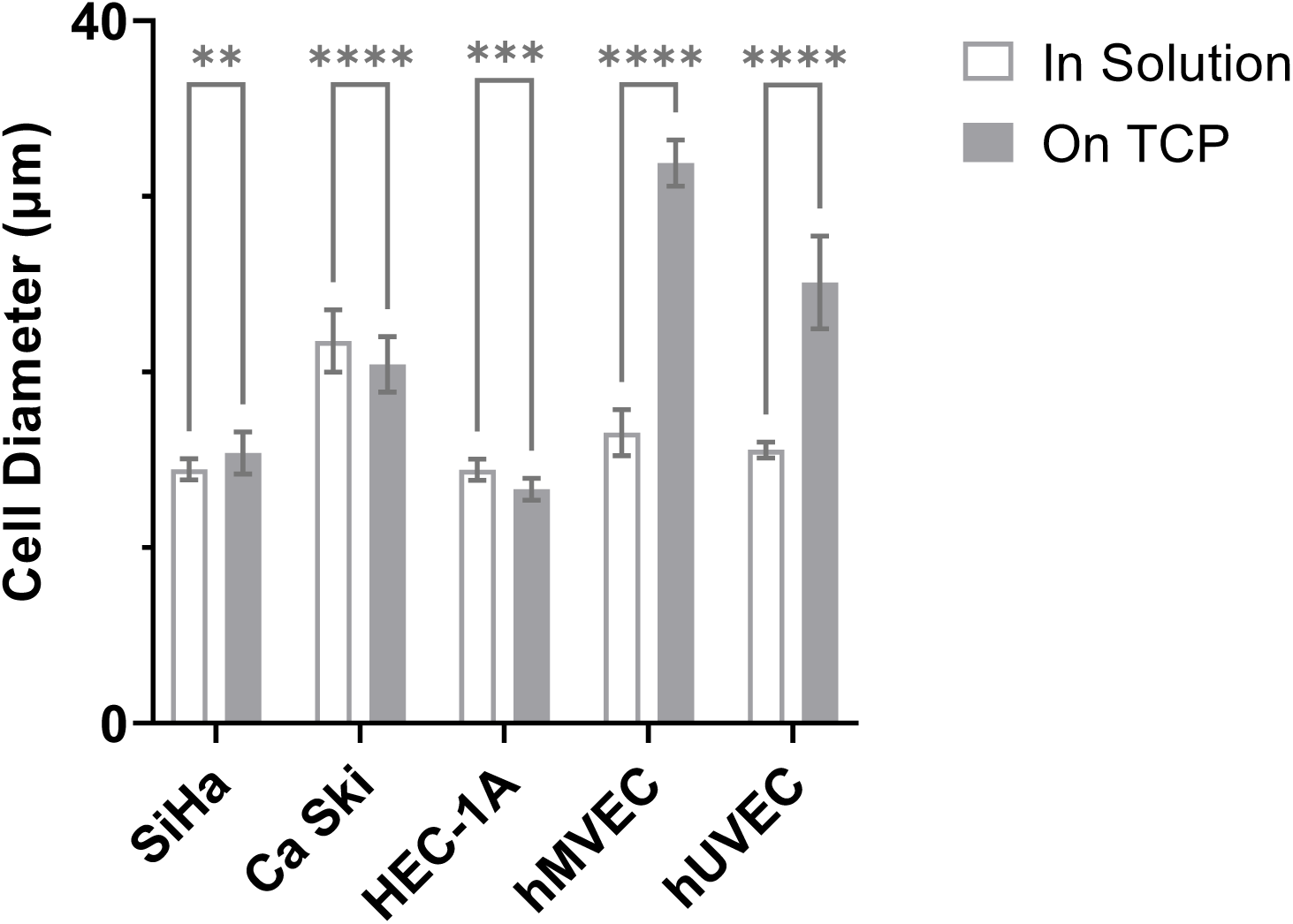
Comparison of cell size in solution and adhered to TCP of cancer cell lines (SiHa, Ca Ski, and HEC-1A) and endothelial cell lines (hMVEC, hUVEC). Statistical analysis completed by 2-way ANOVA with Tukey post-test is included in supplemental data. ns < 0.12, * p <0.05 compared with the mean of each group. Two-way ANOVA with Sidak post-test. Data represent the mean ± SD (30 < n < 120).

There were significant differences in cell diameter for both the cancer and endothelial cell lines in solution compared to on TCP (**Fig. 3**). The SiHa cell diameter in solution was significantly smaller than the diameter measured on TCP, while the diameters for Ca Ski and HEC-1A cells were larger when measured in solution. However, both endothelial cell lines (hMVEC, hUVEC) were found to have a significantly larger diameter when adhered to TCP. Additionally, we calculate the TPR and FPR for single cells dispensed in 96 well plates (**Sup. Table 1**). Considering cell diameter, TPR, and FPR we chose to move forward with a pinch point diameter of 14 µm for all automatic cell dispensing.

### 3.3 Precision, proliferation rate, and viability of MD and AD cancer cell lines

To compare if there were differences between manual and automatically dispensed cells, we evaluated the precision at day 0 and the proliferation rate and viability at day 7. For precision, both dispensing methods for SiHa achieved approximately a precision of one, indicating that the target number of cells per well were equal to the actual number of cells for all cell lines for all dispense methods, (**Fig. 4A**). For Ca Ski the manual dispense cell count was nearly two-fold the target number, while automatic dispensing achieved approximately one. For HEC-1A both manual and automatic dispense achieved an average precision slightly lower than one with automatic dispensing being closer to one than manual dispense. There are no significant differences in the precision measurements between the manually and automatically dispensed replicates for all cell lines, but generally the standard deviation for the automatically dispensed groups is smaller than the standard deviation of the automatically dispensed replicates. For proliferation rate, both dispensing methods yielded a proliferation rate greater than one for SiHa and HEC-1A (**Fig. 4B**). However, for Ca Ski both dispense methods resulted in a mean proliferation rate less than one. There were significant differences in proliferation rate between the manually and automatically dispensed replicates for the SiHa and HEC-1A cell lines, with the mean value of the manually dispensed replicates being greater for all cell lines. For viability, both dispense methods for all cell lines achieved a mean viability greater than 50%, with Ca Ski having the greatest standard deviation for both manually and automatically dispensed cells (**Fig. 4C**).

**Figure 4.**
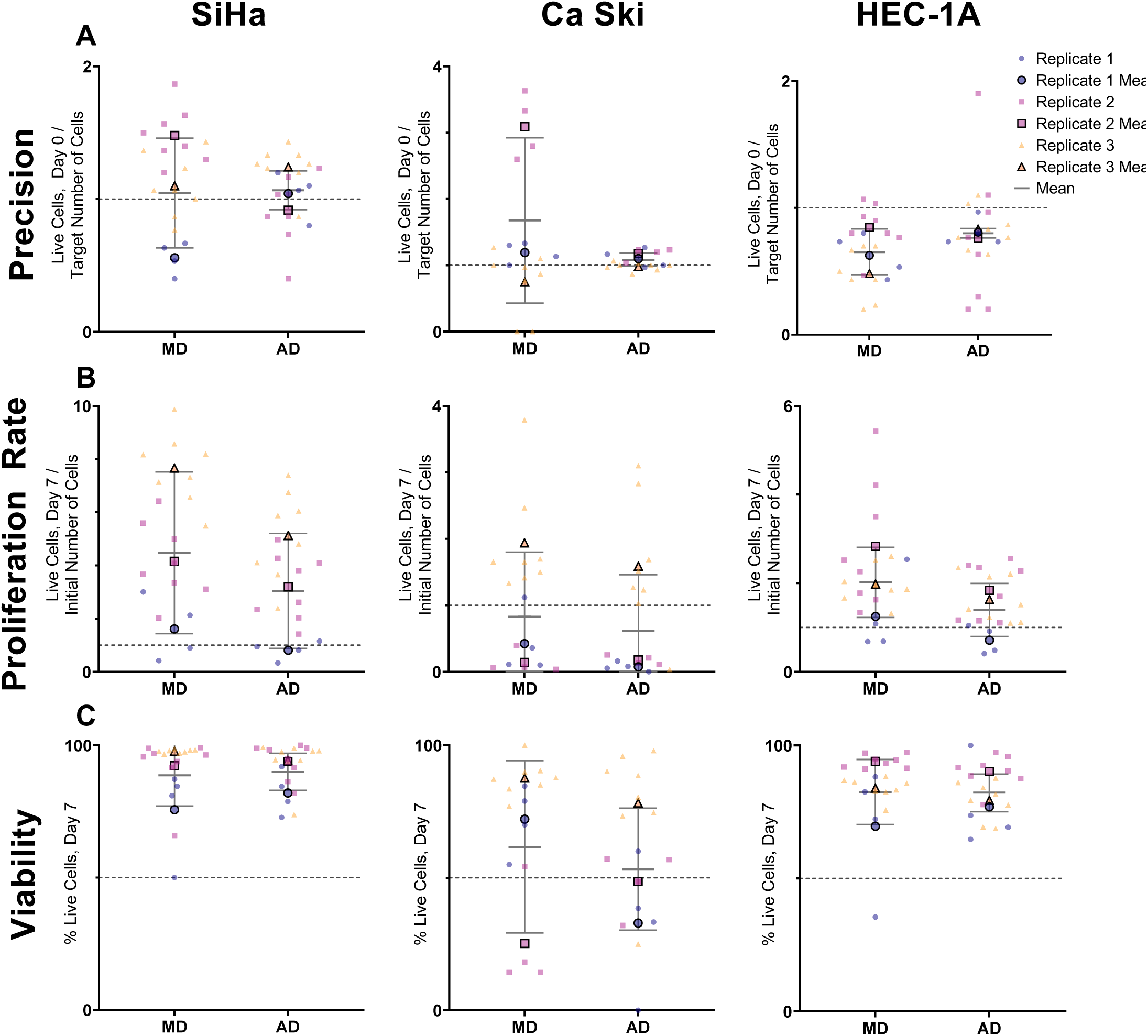
Precision (**A**), proliferation rate (**B**), and viability (**C**) of manual dispense (**MD**) and automatic dispense (**AD**) cancer cell lines (SiHa, Ca Ski, HEC-1A). Cells were dispensed on top of 25 μL of cell specific culture medium on TCP in a 384 well plate. The precision assessment compares the number of live cells MD and AD, evaluated same day of dispense. Cell viability and proliferation evaluated seven days after dispense. Statistical analysis completed by paired t test. ns < 0.12, * p < 0.05. Data represents the mean ± SD (n=4 or 8 per replicate).

### 3.4 Manual vs automatic dispense of cancer cell lines on multi-layer models

To determine if the dispense method affected cell phenotypic dynamics, endothelial cell coverage (**Fig. 5A**), microvessel length (**Fig. 5B**), cancer cell displacement (**Fig. 5C**), and number of cancer cells alive per well (**Fig. 5D**) were measured for both automatically and manually dispensed cancer cells. The only significant differences were cervical cancer cell coverage in the manually dispensed Ca Ski model and a significant increase in the number of cancer cells in the HEC-1A automatically dispensed model (**Fig. 5A**, **Fig. 5D**). In contrast, there were more significant differences observed when the models were formulated with Matrigel (**Fig. 6**). For Matrigel, at least one cell phenotype was significantly different for each cell phenotype.

**Figure 5.**
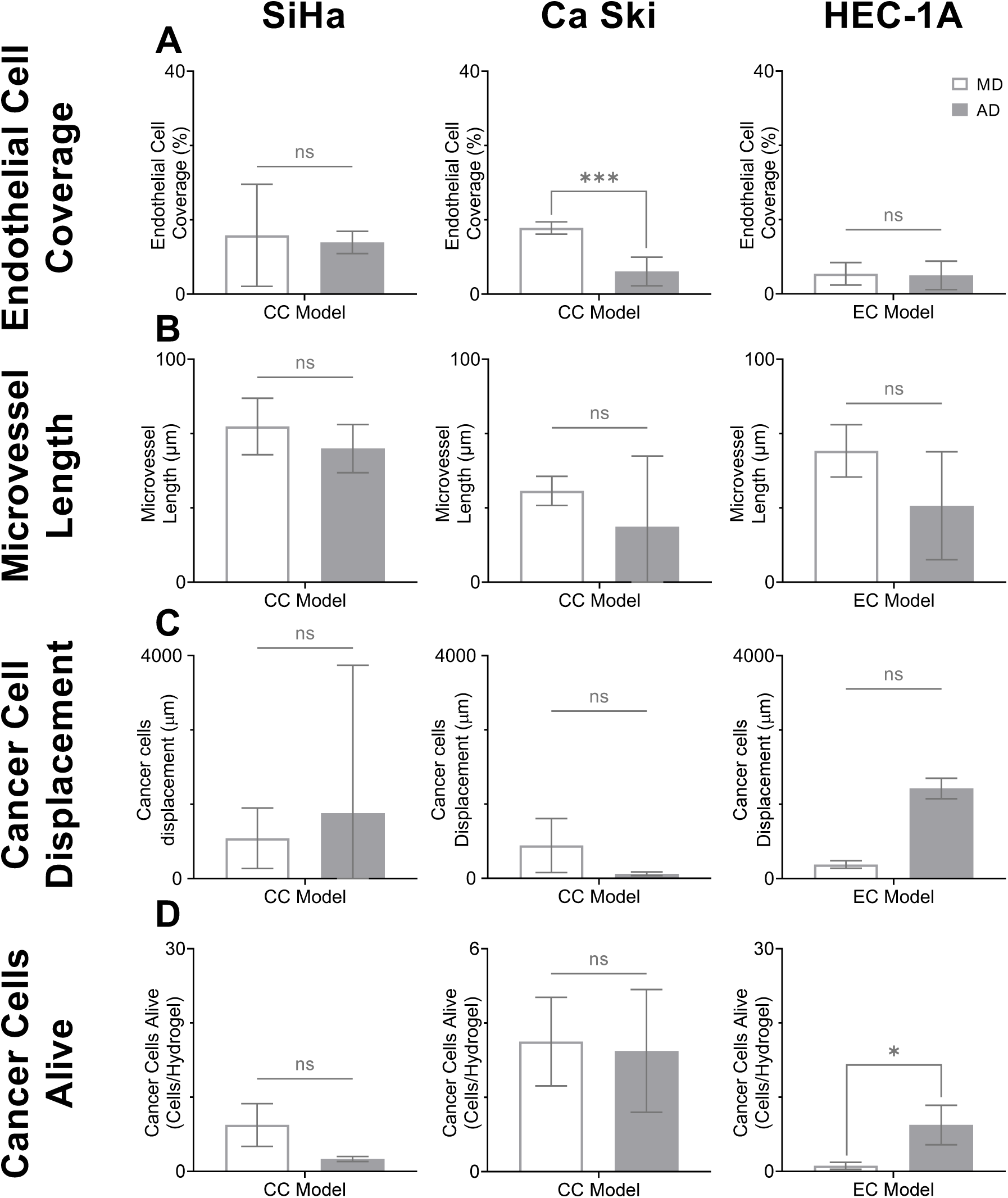
Comparison of manual dispense (**MD**) and automatic dispense (**AD**) in multicellular multilayer models for cervical cancer (**CC**) and endometrial cancer (**EC**). Endothelial cell coverage (**A**), microvessel coverage (**B**), cancer cells displacement (**C**), and number of cancer cells alive (**D**). Endothelial cells (hMVECs) were co-cultured with the cervical cancer cell lines (SiHa, Ca Ski) in the optimized hydrogel model for cervical cancer. Endothelial cells co-cultured with the endometrial cancer cell line (HEC-1A) in the optimized hydrogel model for endometrial cancer. Data represented 24h post culture. Cell viability was asses after 48h of cultured. ns < 0.12, * p <0.033, ** p<0.0021, *** p <0.0002, **** p< 0.0001 compared with the mean of each group. Two-way ANOVA with Sidak post-test. Data represent the mean ± SD (n = 4)

**Figure. 6.**
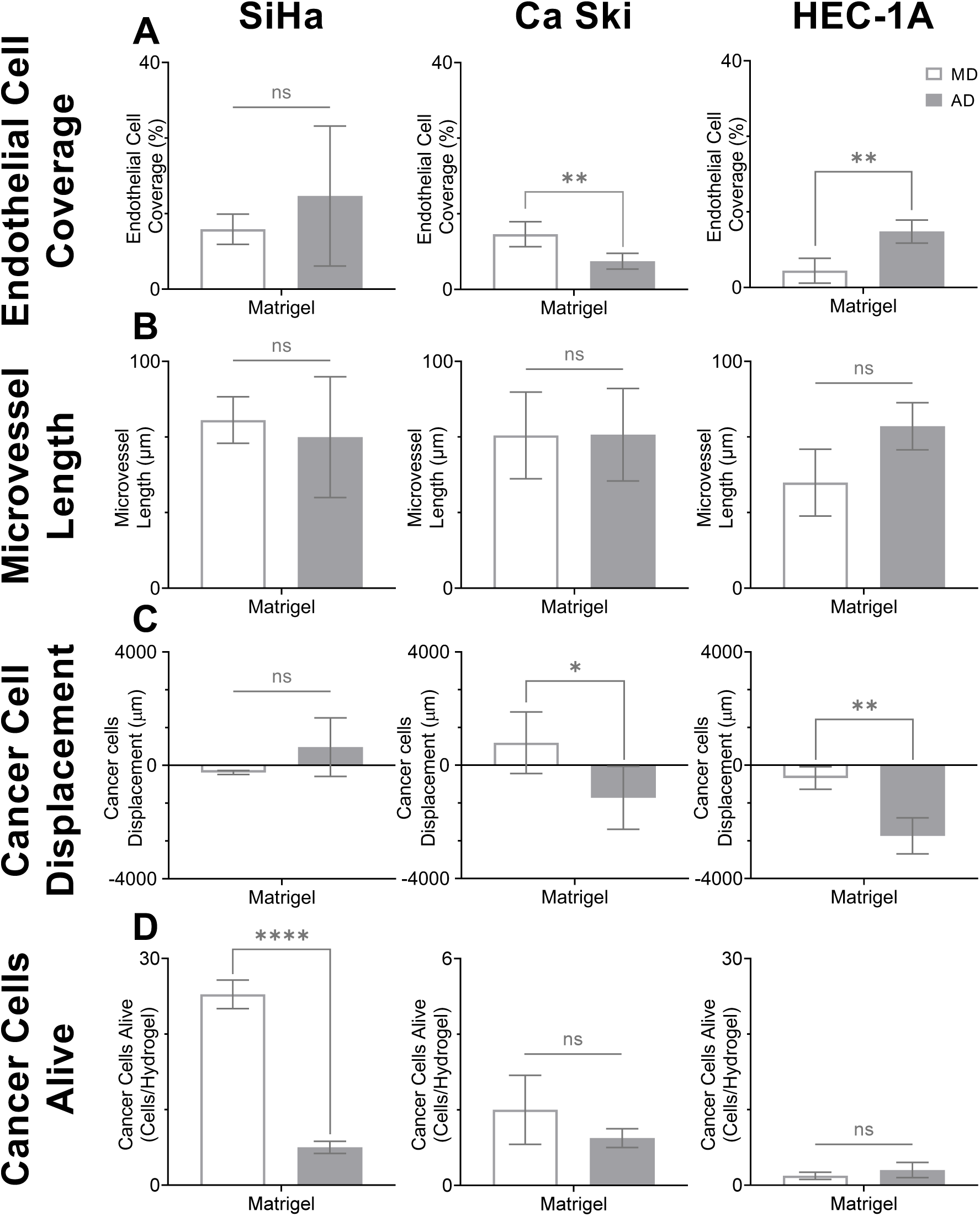
Comparison of manual dispense (**MD**) and automatic dispense (**AD**) in multilayer Matrigel models. Endothelial cell coverage (**A**), microvessel coverage (**B**), cancer cells displacement (**C**), and number of cancer cells alive (**D**). Endothelial cells (hMVECs) were co-cultured with the cervical and endometrial cancer cell lines (SiHa, Ca Ski, HEC-1A) in the Matrigel models. Data represented 24h post culture. Cell viability was asses after 48h of cultured. ns < 0.12, * p <0.033, ** p<0.0021, *** p <0.0002, **** p< 0.0001 compared with the mean of each group. Two-way ANOVA with Sidak post-test. Data represent the mean ± SD (n = 4)

## 4 Discussion

The advancement of precise automated cell dispensing has the potential to enhance reproducibility without changing cell behaviors and signaling. Device limitations come down to the mechanics of dispensing and volume limitations. Both metrics should be properly aligned with research goals. Microliter fluid handlers excel at dispensing precise volumes of reagents rapidly, but lack the ability for precise cell dispensing.^21^ However, microfluidics combined with inkjet technology enables picolitre droplets leading to single cell dispense.^21,22^ Thus, the goal of this work was to evaluate the performance of a single cell dispenser compared to the traditional approach of manual dispensing.

While there are certain applications for dispensing a single cell per well, long term cultures require cells to be dispensed in proximity to other cells to remain viable and proliferate.^25^ Thus to use a single cell dispenser a key first step is to identify a minimum seeding density for each type of cell. When we evaluated cancer cell lines SiHa, Ca Ski, HEC-1A, we found that overall 27 cells per well maintained cell viability and proliferation in a 384 well plate. These values were slightly lower than values previously reported in immortalized and primary cell lines.^26–29^ Too few cells in an individual well and cells demonstrate poor viability and proliferation as cells rely on paracrine and juxtracrine signally to survive.^25^ At the same time, cells seeded at high cell densities maintain high cell viability but exhibit decreased proliferation as confluence can have adverse effects on proliferation.^30–32^ For these reasons all investigations utilize a seeding density of 30 cancer cells per well. When performing single cell dispense with ink-jet technology where the target is multiple cells per well it is pertinent to determine the optimal number of cells to maintain cell behavior and optimize dispensing.

The HP D100 Single Cell Dispenser is an inkjet technology that operates by monitoring the pinch point within a microfluidic channel (**Fig. 6A**). When a cell passes through this the pinch point (funnel), the cell is then flushed into the desired well or unit (**Fig. 6B**). Therefore, the pinch point in the dispenser must be compatible with the cell size. If the pinch point is significantly larger than the cell size, the dispensing may be inaccurate, while a pinch point diameter significantly smaller than the cell diameter has the potential to shear or lyse the cell and alter cell integrity.^22^ We have demonstrated that there are significant differences in cell diameter for cancer cell lines (SiHa, Ca Ski, HEC-1A) and endothelial cell lines (HMVEC, HUVEC) when adhered to TCP compared to cells suspended in media, while not readily reported in literature. As cells are dispensed in solution, it is critical to measure cell diameter that reflects the cell size as it passes through the dispensing mechanisms to avoid adverse effects on the cell.

Once dispensing parameters and cell metrics have been determined, the concern then comes to whether the dispensing mechanism alters cell behavior. Dispensing cells can cause mechanical and morphological changes that can alter function due to the tension, compression, and shear effects that cells are exposed to during the dispensing process.^33^ For monoculture applications, we observed few significant differences in cell function between manually and automatically dispensed cells. We observed increased precision with automatic dispensing while achieving acceptable viability and proliferation. Manual dispensing has been shown to vary greatly not only across handlers but between replicates conducted by just one technician.^34^ However, automated and robotic liquid handling is able to reduce technical variation in dispense, while performing with higher sterility standards.^20,35,36^ With the success of adequate automatic dispensing in monoculture applications, we then deployed the automatic dispensing of cancer cells in co culture applications.

Precision cell dispensing required downsizing of the multilayer multicellular models from our previously reported 96 well plate format to a 384 well plate format.^23^ When automatically dispensing the cancer cells, we found that for our optimized hydrogels there were few significant differences between the phenotypic responses for the manually and automatically dispensed cells. However, we observed more significant differences when our optimized hydrogel was replaced with layers of Matrigel. This is also reflected in the monoculture viability and proliferation investigation of cells that were seeded on Matrigel and could be explained by poor cell adhesion. Matrigel as a platform is less tunable and has been shown to be more genetically different than other ECM hydrogels from human tissues, while showing decreased attachment, proliferation, and differentiation for cells when seed on polystyrene.^37,38^ The ability to automatically dispense the cancer cells on the multilayer multicellular models brings us one step closer to an automatically fabricated model, while the few differences between cell dispensing methods is greatly outweighed by the strides taken towards complete automated dispense.

## Conclusion

We have identified a single cell dispensing method for fabricating 3D *in vitro* multilayer multicellular models with minimal differences between automatic dispensing and traditional dispensing methods without significantly altering cell or model behavior. When determining the feasibility of automatic single cell dispensing on a project-by-project basis is our recommendation to align target dispensing volume with dispenser capabilities, investigate cell size to compliment microfluid restrictions, and monitor any differences between in cell or model behavior between manual and automatic dispensing. Single cell dispensing is recommended for applications where precise cell dispensing is crucial, while bulk cell dispensing is recommended for applications that require high cell seeding density. In conclusion, single cell dispensing can be utilized when developing methods for automatic model fabrication.

## Supplemental

**Sup. Table 1.** Cancer cell lines and endothelial cell line diameter in solution and on TCP.

| Cell Type |  | Cell Size (μm) |  |
| --- | --- | --- | --- |
|  |  | In Suspension | On TCP |
| Cancer Cell Lines | SiHa | 14.5 ± 0.6 | 15.4 ± 1.2 |
|  | Ca Ski | 21.8 ± 1.8 | 20.4 ± 1.6 |
|  | HEC-1A | 14.4 ± 0.6 | 13.3 ± 0.6 |
| Endothelial Cell Lines | HMVEC | 16.5 ± 1.3 | 31.9 ± 3.1 |
|  | HUVEC | 15.7 ± 0.5 | 25.1 ± 2.7 |

**Sup. Table 2.** True positive rate (TPR), false positive rate (FPR) for single cell dispense of cancer and endothelial cell lines for cassettes of 11 μm and 14 μm pinch point.

| Cell Type |  | Metric (%) | Pinch Point Diameter (μm) |  |
| --- | --- | --- | --- | --- |
|  |  |  | 11 | 14 |
| Cancer Cell Lines | SiHa | TPR | 83.61 | 75.00 |
|  |  | FPR | 34.29 | 11.11 |
|  | CaSki | TPR | 60.00 | 76.92 |
|  |  | FPR | 67.21 | 38.60 |
|  | HEC-1A | TPR | 87.50 | 91.38 |
|  |  | FPR | 37.50 | 28.95 |
| Endothelial Cell Lines | hMVEC | TPR | 68.57 | 84.09 |
|  |  | FPR | 38.30 | 36.54 |
|  | hUVEC | TPR | 56.25 | 80.00 |
|  |  | FPR | 67.50 | 66.67 |

**Sup. Fig. 1.**
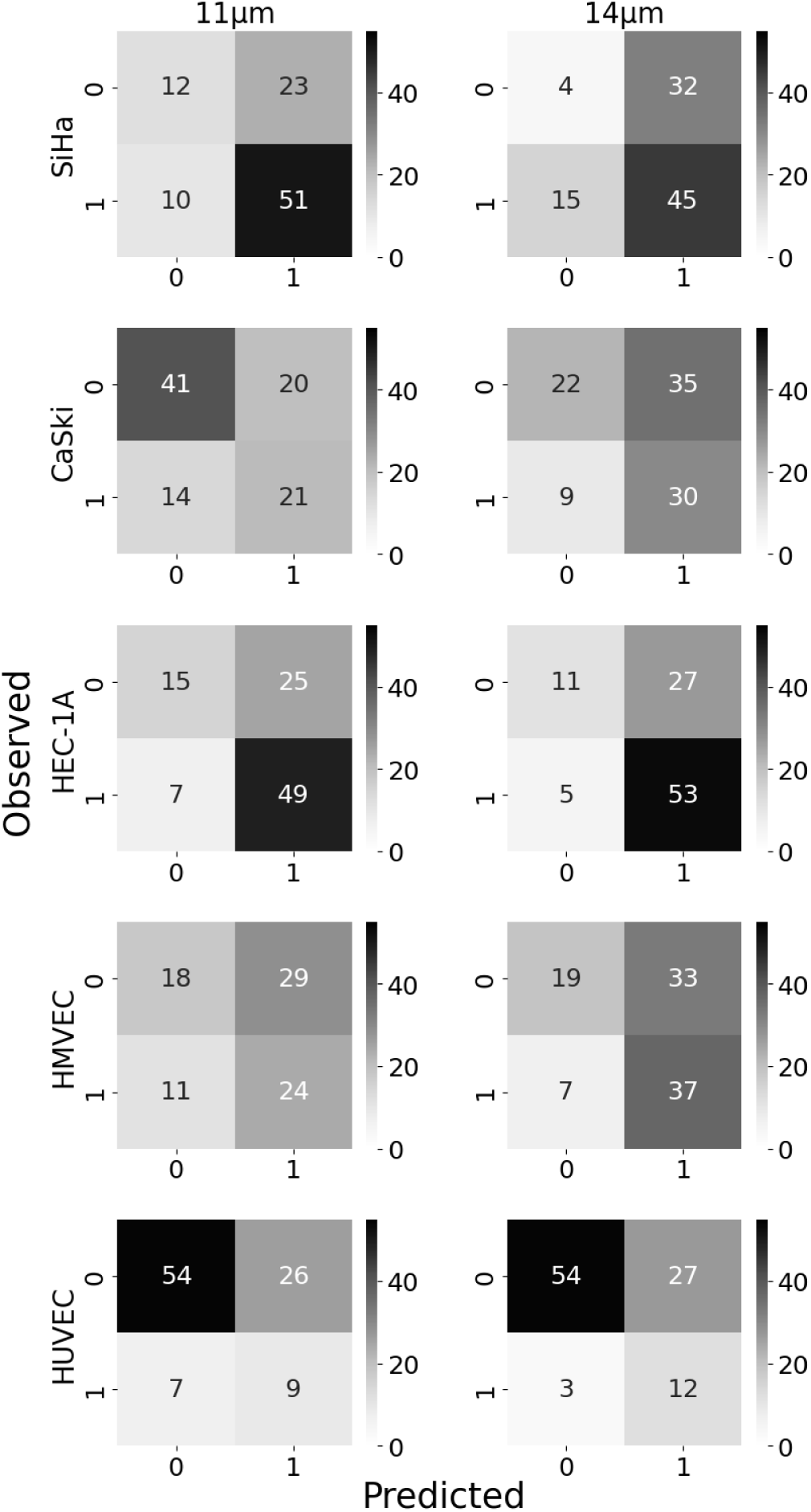
Comparison of cassette pinch point diameter (11µm and 14 µm) on for the cancer cell lines (SiHa, Ca Ski, and HEC-1A) and the endothelial cell lines (hMVEC and hUVEC). The cell lines were automatically dispensed on top of 10 µL of culture media in a 96 well plate. Results are presented in a confusion matrix where predicted data represents dispenser detection of single cell wells (1) and non-single cell wells (0) and observed single cell wells and non-single cell wells represents image collected data. Wells predicted and observed to be a single cell well represent a true positive (TP) value, while wells predicted and observed to be non-single cell wells represent true negative (TN) value. Wells predicted to be a single cell well but observed to be non-single cell wells are indicative of a false positive (FP), and wells predicted to be a non-single cell wells but observed to be a single cell wells are indicative of a false negative (FN) value.

